# Distributional coding of associative learning within projection-defined populations of midbrain dopamine neurons

**DOI:** 10.1101/2022.07.18.500429

**Authors:** Riccardo Avvisati, Anna-Kristin Kaufmann, Callum J. Young, Gabriella E. Portlock, Sophie Cancemi, Rui Ponte Costa, Peter J. Magill, Paul D. Dodson

**Author notes:** These authors contributed equally to the work (Riccardo Avvisati, Anna-Kristin Kaufmann).

## Abstract

Midbrain dopamine neurons are thought to play key roles in learning by conveying the difference between expected and actual outcomes. While this teaching signal is often considered to be uniform, recent evidence instead supports diversity in dopamine signaling. However, it remains poorly understood how heterogeneous signals might be organized to facilitate the role of downstream circuits mediating distinct aspects of behavior. Here we investigated the organizational logic of dopaminergic signaling by recording and labeling individual midbrain dopamine neurons during associative behavior. We defined combinations of protein expression and cellular localization to sort recorded neurons according to the striatal regions they innervate. Our findings show that reward information and task variables are not only heterogeneously encoded, with multiplexing, but also differentially distributed across populations of dopamine neurons projecting to different regions of striatum. These data, supported by computational modelling, indicate that such distributional coding can maximize dynamic range and tailor dopamine signals to facilitate the specialized roles of different striatal regions.

## INTRODUCTION

Learning to anticipate positive or negative consequences from environmental cues is essential for survival. Midbrain dopamine neurons are thought to play key roles in this process by signaling the difference between expected and actual outcomes (reward prediction error; RPE) (Schultz et al., 1997). Such a fundamental teaching signal has traditionally been thought to necessitate a uniform message (Eshel et al., 2016; Schultz, 2016a, 2016b). However, recent work has reve that dopamine neurons and the signals they convey show considerable heterogeneity. For example differences in dopamine release and dopamine axon activity have been observed in different striatal regions (Brown et al., 2011; de Jong et al., 2019; Parker et al., 2016; Saddoris et al., 2015; Tsutsui-Kimura et al., 2020; Yuan et al., 2019) with temporally-distinct activity patterns recorded across dorsal striatum (Hamid et al., 2021). Furthermore, dopamine neurons seem to signal more than just reward; encoding movement onset, movement kinematics, and multiple variables involved in decision-making (Barter et al., 2015; Coddington and Dudman, 2018; Dodson et al., 2016; Engelhard et al., 2019; Howe and Dombeck, 2016; Hughes et al., 2020; Kremer et al., 2020; da Silva et al., 2018). Dopamine neurons are also molecularly, physiologically, and anatomically diverse; single-cell RNA sequencing has identified multiple groups of midbrain dopamine neurons that can be distinguished by the combinatorial expression of different molecular markers (Hook et al., 2018; La Manno et al., 2016; Poulin et al., 2014, 2020; Saunders et al., 2018a; Tiklová et al., 2019), and there is evidence that dopamine neurons exhibit differences in ion channels, other protein expression, firing properties, and input-output connectivity (Farassat et al., 2019; Lammel et al., 2008, 2011; Lerner et al., 2015; La Manno et al., 2016; Morales and Margolis, 2017; Poulin et al., 2014, 2020; Roeper, 2013; Schiemann et al., 2012; Watabe-Uchida et al., 2012). Together, these findings suggest that there might be specialized midbrain populations that each convey different information to discrete areas of striatum. However, it is not yet clear how heterogenous signals might instruct different striatal regions that each play diverse and complimentary functional/behavioral roles. For example, the dorsolateral striatum (DLS) is thought to subserve sensorimotor functions in stimulus-response associations and habits, whereas the dorsomedial striatum (DMS) is involved in response-outcome associations for goal-directed actions (Balleine and O’Doherty, 2010). In further contrast, the ventrolateral striatum (VLS) is thought to be important for motivation (Tsutsui-Kimura et al., 2017), whereas the core of the nucleus accumbens (NAc) has been ascribed roles in outcome-evaluation (Saddoris et al., 2015; Saunders et al., 2018b).

To define the organizational principles underlying heterogeneous dopamine signaling, we recorded and molecularly-identified individual dopamine neurons in mice during Pavlovian conditioned behavior. We defined combinations of protein expression and cellular localization to sort recorded neurons according to the striatal regions they innervate. We find that there is no broad spatial organization of encoding in midbrain, but instead we show that neurons projecting to the same target regions are more homogenous in their firing patterns and that these populations are distinct from those projecting to other parts of striatum. Temporal difference modelling predicts that distributional coding within projection-defined populations can tailor them to support different aspects of associative learning.

## RESULTS

### Dopamine Neurons Heterogeneously Encode Reward Behavior

We trained head-fixed mice in a Pavlovian conditioning paradigm in which an auditory cue (4 kHz tone, 1 s) signaled reward delivery after a fixed delay (2 s from cue onset). Mice rapidly learned to associate the cue with reward as indicated by anticipatory licking (Figure 1A). To investigate the firing of different dopamine neurons during the task, we extracellularly recorded individual neurons and then juxtacellularly labeled them to precisely determine their location and confirm they were dopaminergic (Figure 1B). The majority of dopamine neurons altered their firing rate at the onset of the cue and/or reward (Figures 1B and 1C). However, while any changes at reward tended to be increases in firing, changes at cue onset were a mix of increases and decreases in rate. Firing at cue and reward may reflect the encoding of reward prediction (Schultz, 2007), or alternatively, encoding of actions with which to obtain reward (Barter et al., 2015; Coddington and Dudman, 2018, 2019; Hughes et al., 2020). To investigate whether changes in firing rate correlated with cue, reward, licking, or other body movements, we used a general linear model (GLM) (Figures 1D, 1E, and S1). We found that many neurons encoded reward (26 of 66) and/or cue (20 of 66). However, we also found that a significant number of neurons encoded aspects of the task which were not obvious from the peristimulus time histogram including licking (16 of 66) or movement (10 of 66). A considerable proportion of neurons (19 of 66) did not significantly encode (p > 0.01) any of the classifiers we examined (Figures 1D and 1E), which suggests that there may be populations of dopamine neurons that encode other facets of behavior. Interestingly, many neurons encoded more than one task variable (21 of the 47 neurons encoding task variables), concordant with the idea that dopamine neurons multiplex signals (Engelhard et al., 2019; Kremer et al., 2020). These data suggest that there is not a uniform signal across midbrain dopamine neurons, but instead support a framework of heterogeneity with some neurons encoding single variables and others multiplexing signals to encode several distinct aspects of behavior (Figure 1E).

**Figure 1.**
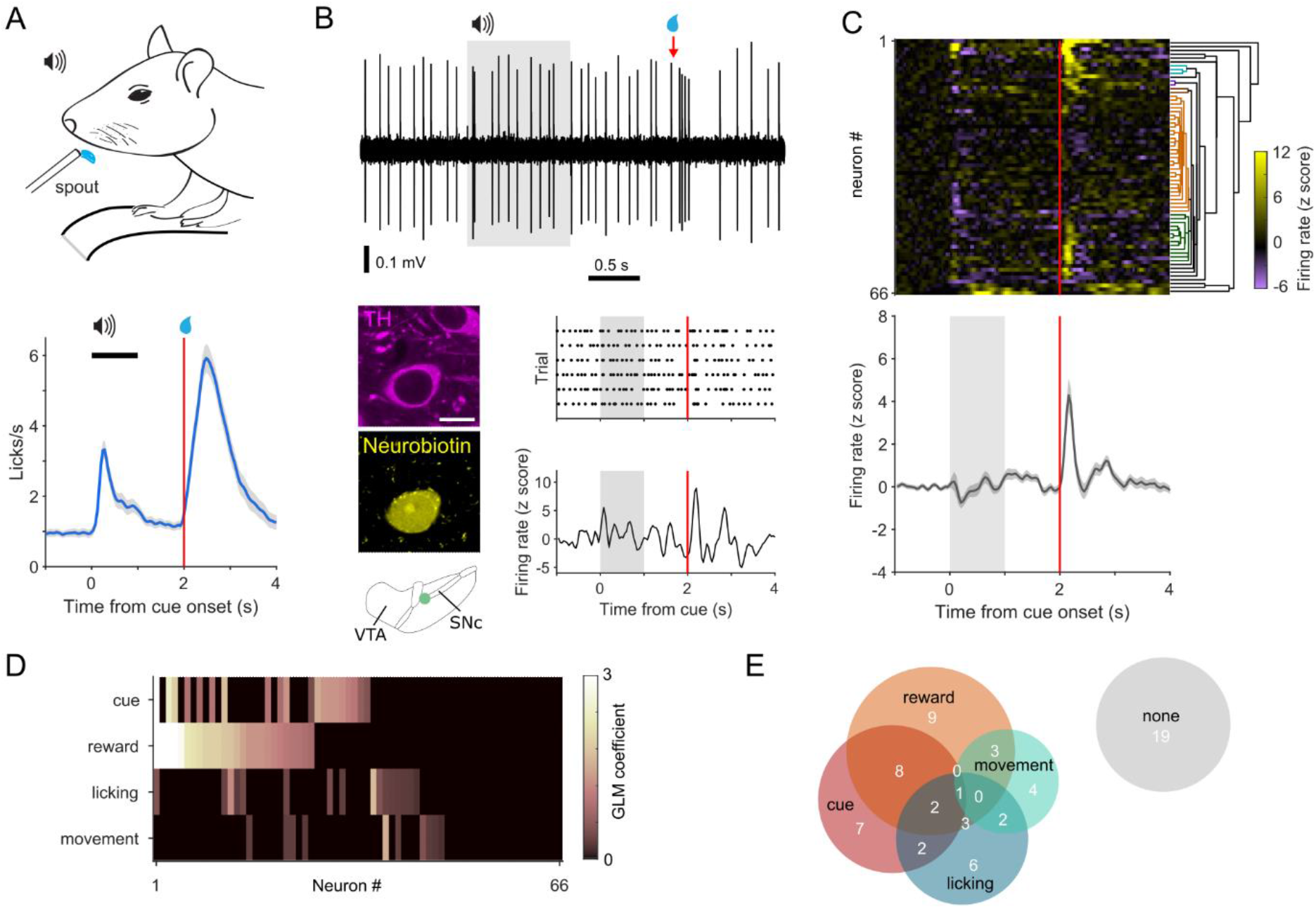
Dopamine neurons heterogeneously encode reward and predictive cues. (A) Head-fixed mice, positioned on a treadmill, were trained to associate a 4 kHz auditory tone (cue) with reward delivery. After training, mice show robust anticipatory licking after the cue and licking to receive reward. (B) Extracellular recording (top) and corresponding peri-stimulus time histogram (PSTH; lower right) from an individual dopamine neuron during Pavlovian conditioned behavior. Grey shading indicates cue duration, red line indicates reward delivery. Neurons were juxtacellularly labeled with neurobiotin and tested for immunoreactivity to tyrosine hydroxylase (TH: lower left; scale = 10 μm) to confirm their dopaminergic identity and localization in the midbrain. The schematic depicts the location of the neuron in the dopaminergic midbrain. (C) Z-scored PSTH of individual responses from identified dopamine neurons (rows) grouped by hierarchical clustering (top), and mean response (bottom). (D) Classifiers which correlate with changes in firing rate for each neuron, determined by a general linear model (GLM). (E) GLM output suggests that neurons can multiplex signals by encoding multiple aspects of behavior (numbers indicate the proportion of neurons).

### Encoding is Not Clearly Defined by Anatomical Location

What might underlie the heterogeneity we observe in reward-signaling? Given the different roles ascribed to the ventral tegmental area (VTA) and the substantia nigra pars compacta (SNc), one might expect that neurons in these regions would encode the task differently (Collins and Saunders, 2020; Cox and Witten, 2019; Lerner et al., 2021). To investigate this possibility, we divided neurons into those located in the VTA (n = 31) and SNc (n = 35) and compared their responses during the task. Despite previous observations of differences in encoding of spontaneous body movements by VTA and SNc neurons (Dodson et al., 2016), the average responses of these two groups during Pavlovian conditioned behavior were nearly identical, and we observed similar clustering of responses (Figures 2A and 2B). The comparable proportions of neurons in each region encoding cue, reward, licking, and movement (Figures 2A and 2B), suggest that the two regions encode the task in a similar manner. However, it is possible that such a blunt subdivision may mask finer spatial organization. We therefore considered whether encoding was organized in a way that neurons in close proximity signaled similar task variables. To probe this possibility, we plotted the dominant task feature encoded by each neuron in Cartesian space (Figures 3A and 3B). We found that all task variables were represented across the anteroposterior and mediolateral extent, arguing against a precise focal encoding of task variables in different regions of the dopaminergic midbrain.

**Figure 2.**
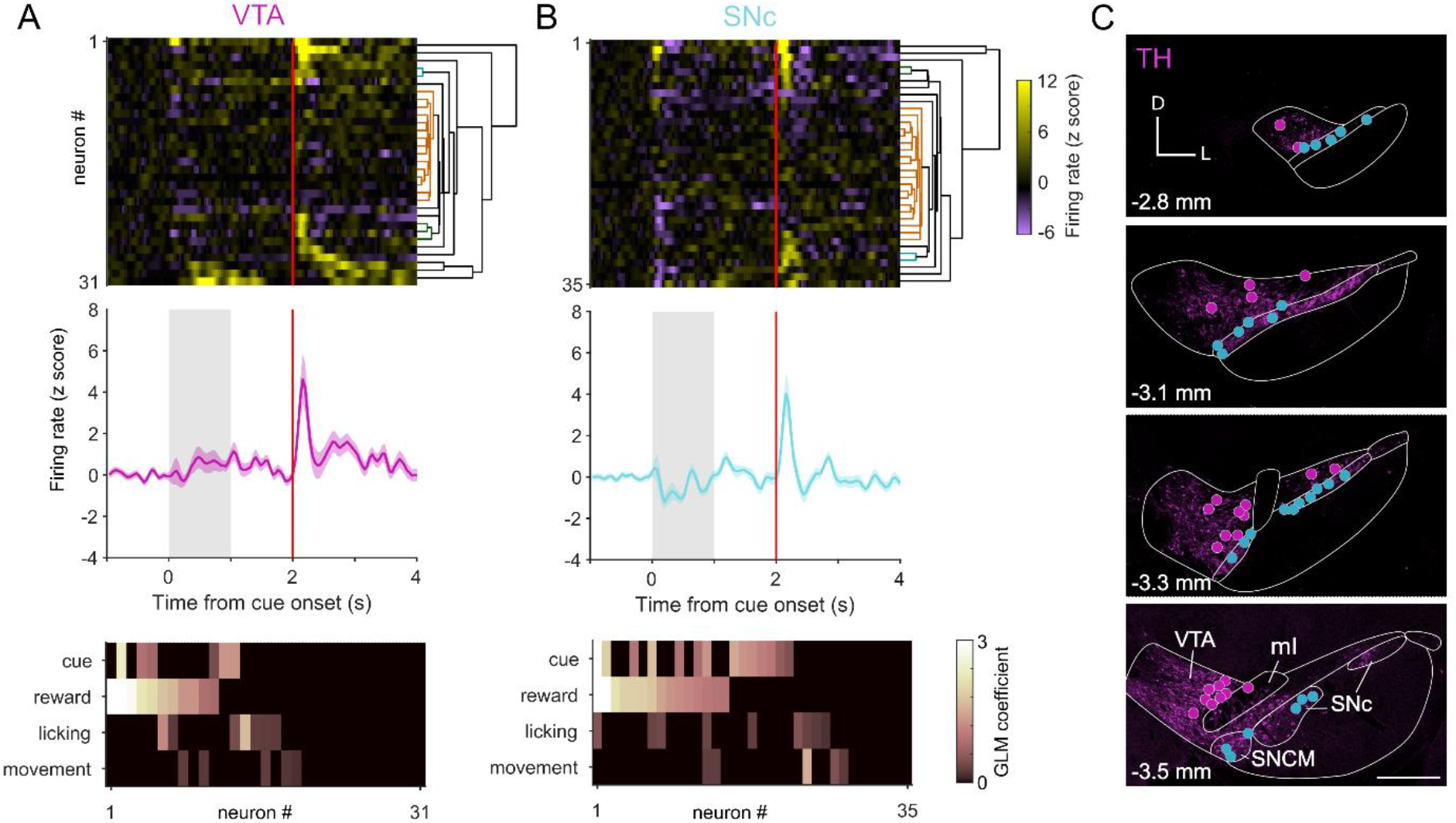
Anatomical subgroups do not account for response heterogeneity. (A) VTA neurons heterogeneously encoded aspects of Pavlovian conditioned behavior (top) and firing could be classified as encoding multiple task variables by GLM (bottom). (B) SNc neurons exhibit a similar degree of heterogeneity in firing responses. (C) Schematic of the locations of recorded and juxtacellularly labeled neurons (VTA neurons magenta, SNc cyan) overlayed on example images of tyrosine hydroxylase (TH) immunofluorescence at four anteroposterior positions 2.8 to 3.5 mm from bregma. D=dorsal; L=lateral; VTA=ventral tegmental area; SNc=substantia nigra pars compacta; ml=medial lemniscus; SNCM=medial part of SNc. Scale = 500 μm.

**Figure 3.**
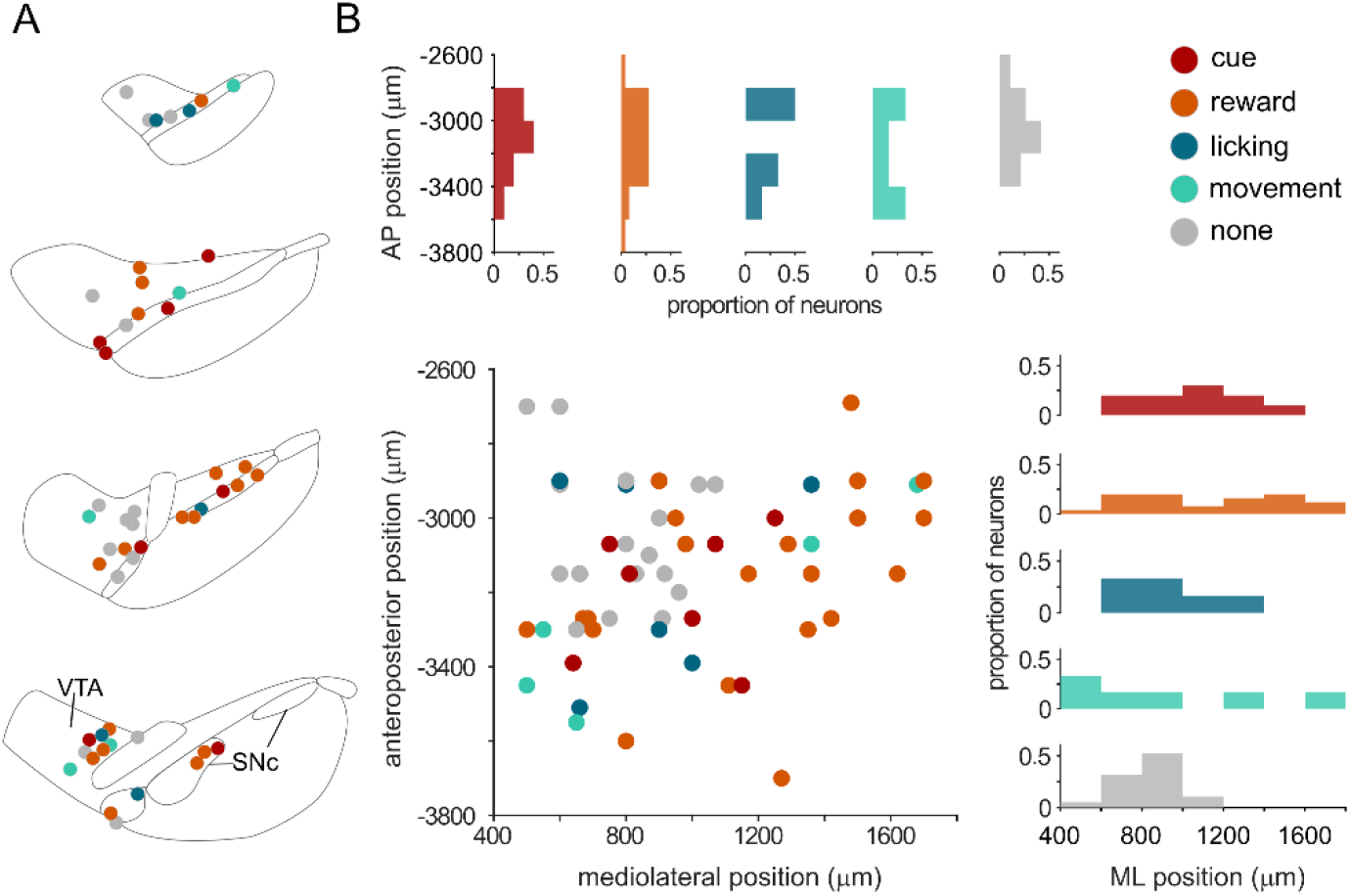
Encoding of task variables is not focally organized. (A) Schematic of the dominant variable encoded by each dopamine neuron according to their location in the midbrain. (B) The dominant variable plotted according to the neurons anteroposterior (AP) or mediolateral (ML) position. The proportion of neurons encoding each dominant variable is shown in the AP (top) and ML plane (right)

### Neurons Projecting to Different Striatal Regions Express Alternate Combinations of Proteins

If encoding is not spatially organized in the midbrain, is there another structural principle underlying response heterogeneity? Emerging evidence suggests that dopamine neurons projecting to particular target regions may differently encode task variables (de Jong et al., 2019; Lerner et al., 2015; Parker et al., 2016; Saunders et al., 2018b). We therefore considered whether distinct midbrain-striatal circuits may account for some of the heterogeneity we observe. To ascribe projection identity to dopamine neurons, but avoid potentially altering dopamine neuron signaling by expressing fluorescent reporter proteins (Farassat et al., 2019; Schiemann et al., 2012), we used a method that took advantage of the known heterogeneity in the expression of endogenous proteins (Poulin et al., 2020). We considered three proteins (Aldh1a1, Sox6, and calbindin) that exhibit different expression patterns across the dopaminergic midbrain and can be robustly detected by commercially available antibodies. To identify the axonal projection targets of dopamine neurons, we injected retrograde tracer (cholera toxin B subunit; CTB) into the dorsomedial striatum (DMS), dorsolateral striatum (DLS), ventrolateral striatum (VLS), or nucleus accumbens core (NAc core) and then examined the marker-expression in retrogradely-labeled midbrain dopamine neurons using immunohistochemistry. We found that most dopamine neurons projecting to DLS expressed Sox6 and Aldh1a1, but not calbindin, and were located in SNc (Figures 4A – 4D), whereas those projecting to NAc core were located in VTA and had the opposite expression pattern (i.e. calbindin, but not Aldh1a1 or Sox6; Figure 4A – 4C). The marker expression of neurons projecting to DMS was less binary, with moderately prevalent expression of Aldh1a1 and Sox6, and rare expression of calbindin (Figure 4C). However, we found that these neurons are tightly clustered in the medial part of SNc (SNCM; Figure 4B) (Farassat et al., 2019). Therefore, by combining the soma location with the expression of Aldh1a1 and Sox6 we could distinguish DMS-projecting dopamine neurons. Similarly, we identified a population of VLS-projecting neurons, localized to the parabrachial pigmented area of the VTA (PBP) and SNc (Farassat et al., 2019), which expressed Sox6 but not Aldh1a1 (Figure 4B and 4C). Thus, by using a combination of anatomical location and marker expression, we could identify four distinct dopamine neuron populations that each innervated a discrete area of striatum (Figure 4E).

**Figure 4.**
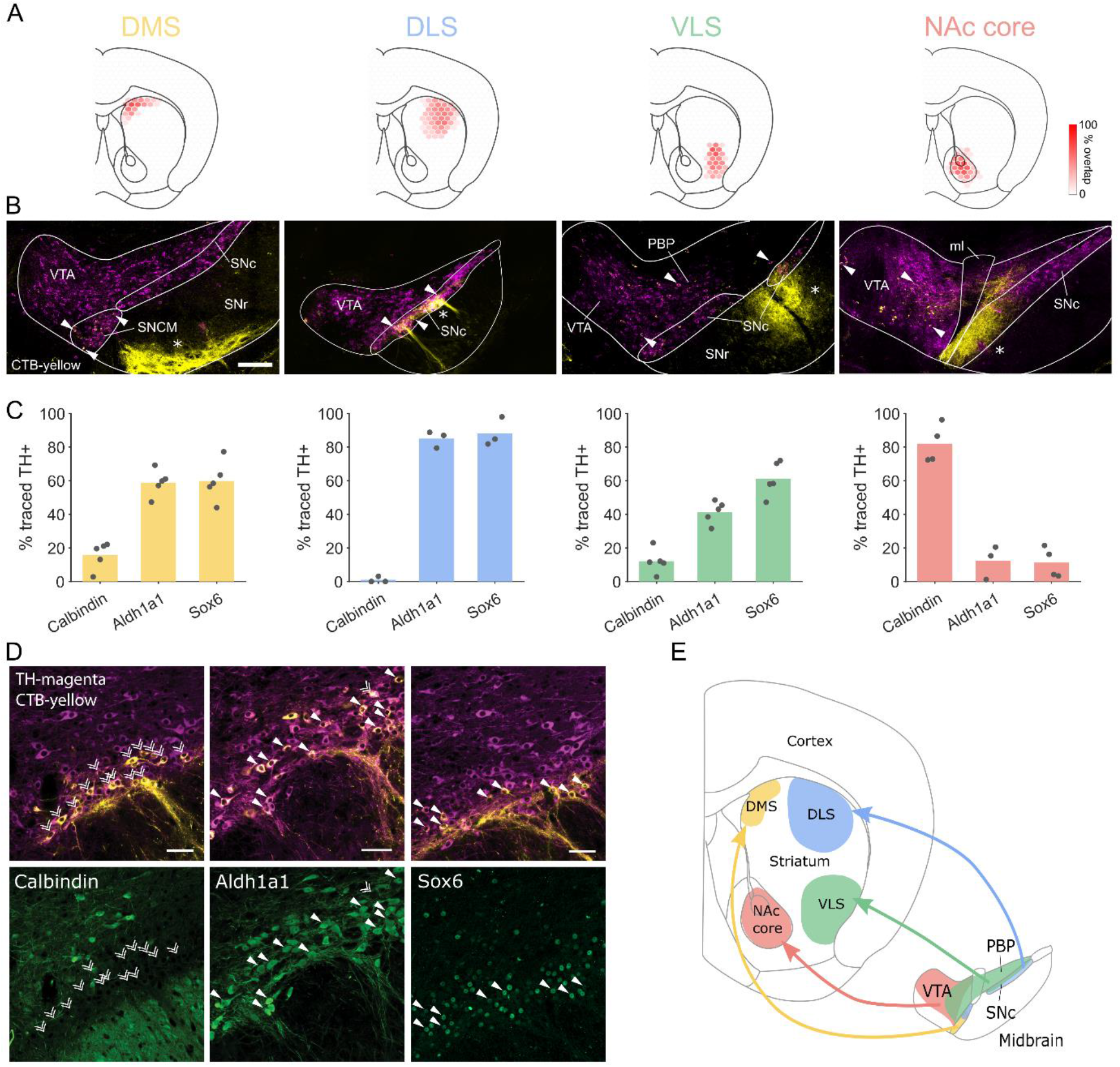
Differential marker expression in populations projecting to different parts of striatum. (A) Schematics showing the extent of retrograde tracer injections into different striatal regions (DMS, dorsomedial striatum; VLS, ventrolateral striatum; NAc core, core of the nucleus accumbens). Colored hexagons represent overlap of CTB injections between animals. (B) Representative midbrain images following CTB injection into different striatal regions. Selected retrogradely traced dopamine neurons (filled arrowheads) illustrate the region of the dopaminergic midbrain which projects to each striatal target. Anterogradely traced fibers (asterisks) from striatal projection neurons are also visible in the SNc and SNr. VTA = ventral tegmental area; SNc = substantia nigra pars compacta; SNr = substantia nigra pars reticulata; ml = medial lemniscus; SNCM=medial part of SNc; PBP=parabrachial pigmented area of the VTA. Scale = 100 μm. (C) Cell counting of CTB positive dopaminergic neurons revealed the percentage of neurons which express each marker. (D) Retrograde tracing with cholera toxin B (CTB) injected into the dorsolateral striatum (DLS) identified SNc dopamine neurons (positive for tyrosine hydroxylase, TH; top panels) which innervate this region. The majority of these neurons expressed Aldh1a1 and Sox6, but not calbindin. Filled arrowheads represent marker expression, double arrowheads, lack of expression. Scale = 50 μm (E) Schematic illustrating the innervation of different striatal regions by neurons in different parts of the dopaminergic midbrain.

### Neurons Projecting to Different Striatal Regions Encode Distinct Aspects of Behavior

Having established criteria that delineated neurons projecting to each of the four striatal target regions, we grouped recorded neurons that matched these criteria and found clear differences in firing pattern between groups (Figures 5A and 5B). On average, putative DLS-projecting and VLS-projecting neurons did not change their firing upon cue presentation (despite the anticipatory licking suggesting that the mice registered and learned the predictive value of the cue), but robustly increased their firing shortly after reward presentation (Figure 5A). In contrast, DMS-projecting neurons exhibited a transient decrease in firing at cue onset and showed no increase in firing at reward presentation (Figure 5A). Putative NAc core-projecting neurons also showed a distinct response, increasing their firing at both cue and reward. However, rather than being time-locked to the onset of these events, these increases in firing were delayed by a few hundred milliseconds, coinciding with periods when the mice were licking. To examine whether these projection-defined groups can account for the heterogeneity observed across all dopamine neurons we performed hierarchical clustering. Clusters were enriched with neurons from a given population, but did not solely contain one population (Figure 5C). For example, 83% of neurons in one cluster were putative VLS-projecting neurons but the remaining VLS-projecting neurons were ascribed to other clusters. It has recently been observed in anaesthetized mice that neurons projecting to different regions have different firing properties (Farassat et al., 2019). We therefore tested whether the populations we identified had distinct firing in awake mice at rest (i.e. during the ITI, outside of engagement with the task). We found that DMS, DLS, and VLS shared similar properties, whereas the NAc core-projecting population had significantly slower firing rates (p < 0.05) and they exhibited longer pauses than other populations (p < 0.01) (Figure S2).

**Figure 5.**
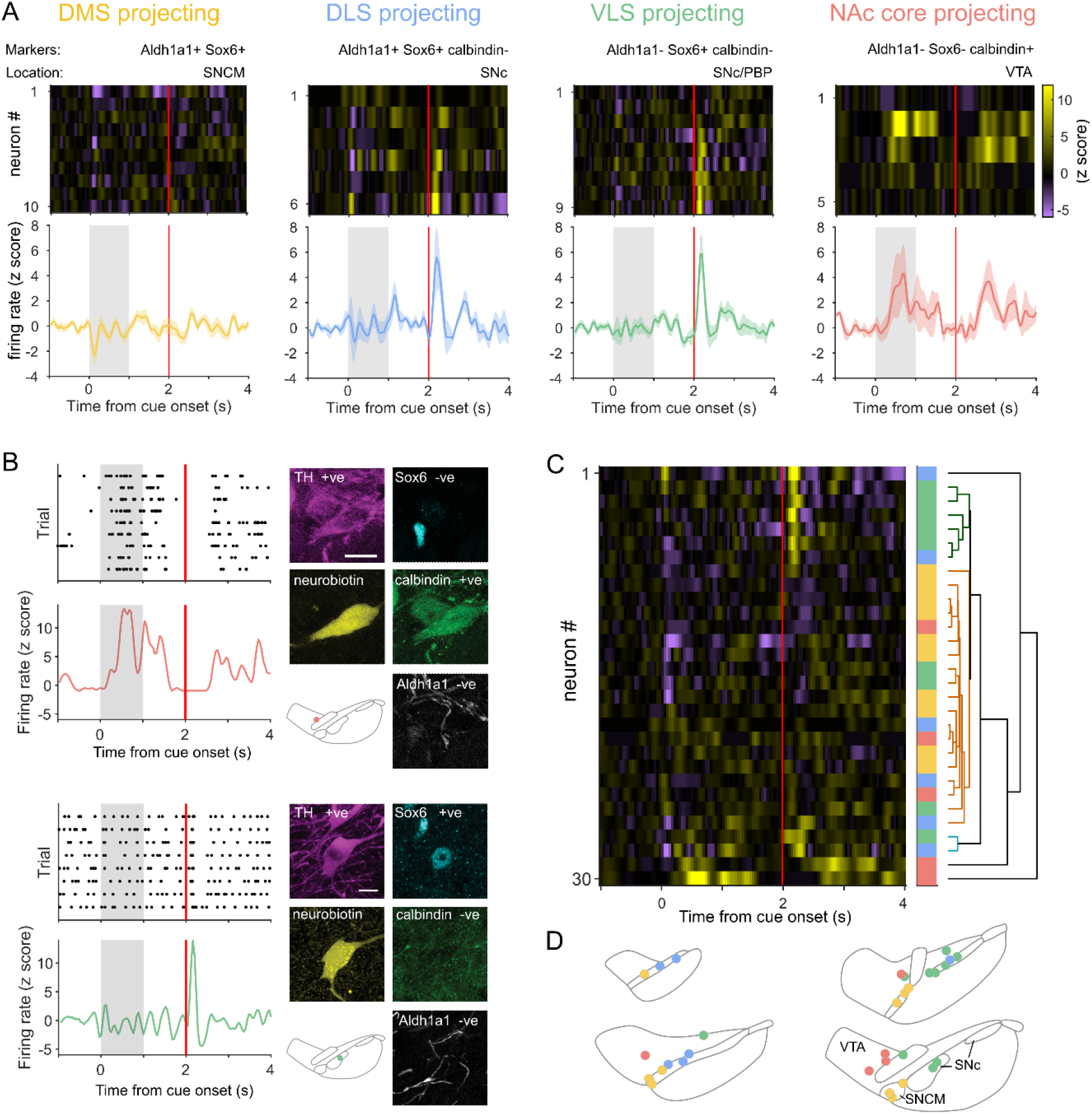
Neurons projecting to different striatal regions exhibit distinct responses. (A) Mean PSTH and individual heatmap plots for neurons putatively projecting to different striatal regions according to classifying marker combinations and location (top). (B) PSTH and marker expression of an example putative NAc core-projecting VTA neuron (top) and VLS-projecting SNc neuron (bottom). Scale = 10 μm (C) Hierarchically clustered individual responses (colored blocks denote putative projection target). (D) Schematic of location of recorded neurons color coded by their putative projection-targets. Neurons in grey did not match the delineating criteria.

Dopamine neurons are considered to encode the difference between actual and expected reward, as such, a perfectly predicted reward should result in no change in firing at reward delivery (Schultz et al., 1997). While DMS- and NAc core-projecting populations exhibited this expected response, many DLS- or VLS-projecting neurons increased their firing at reward delivery (Figure 5A). To probe this further, we plotted the normalized firing at reward for each population. We observed that DLS- and VLS-projecting populations had a broader range of responses suggesting that some neurons were overestimating and others underestimating future reward (Figure 6A). Recent work has suggested that encoding optimism as a probability distribution across dopamine neurons may confer advantage to reward learning (Dabney et al., 2020). Our data suggest that such distributional coding may differ in populations projecting to different parts of striatum in both the width of the distribution and the optimistic skew (Figure 6A). To investigate the effect that different distributions might have on reward learning, we used a simple temporal difference (TD) model where an agent learns state-value associations as it transitions through a ‘Pavlovian conditioning’ environment. Projection-specific agents were generated by biasing reward prediction errors by a term sampled from distributions that approximated experimental data (Figure 6B). To probe whether different distributions confer a general advantage for learning, we tested the accuracy of value estimations made by each projection-defined agent using a wider range of rewards (Figure 6C; reward range 0 – 20). Some striatal regions are thought to be important for state-value associations whereas others are thought to have a role in motivational control (Bromberg-Martin et al., 2010; Matsumoto and Hikosaka, 2009; Saddoris et al., 2015; Schultz, 2007; Tsutsui-Kimura et al., 2017). To examine the accuracy of projection-defined agents in state-value association, we used a ‘familiar’ range of rewards, close to the center of the training range (3.5 – 6.5) whereas to examine motivational control, we considered the performance of agents when rewards are slightly larger than average (60 – 75% of reward range). We reasoned that if a reward is large enough, motivation to obtain the reward should naturally be high, whereas when rewards are just slightly larger than normal, judgement is required to determine whether a reward is worth expending effort to attain. We found that DMS- and NAc core-projecting agents made significantly more accurate state-value associations (as might be expected from their narrow distribution coding) and DLS-projecting agents performed significantly worse than all other groups (Figure 6D). However, DLS- and VLS-projecting agents were considerably more accurate than DMS- and NAc-core agents in the ‘motivation range’, with VLS-projecting agents exhibiting the best performance (Figure 6E). Taken together, this suggests that populations projecting to medial regions of the striatum (DMS and NAc core) convey dopamine signals that may be tuned to support state-value learning, whereas populations projecting to lateral regions (VLS and DLS) might be better suited to providing information to support effortful action.

**Figure 6.**
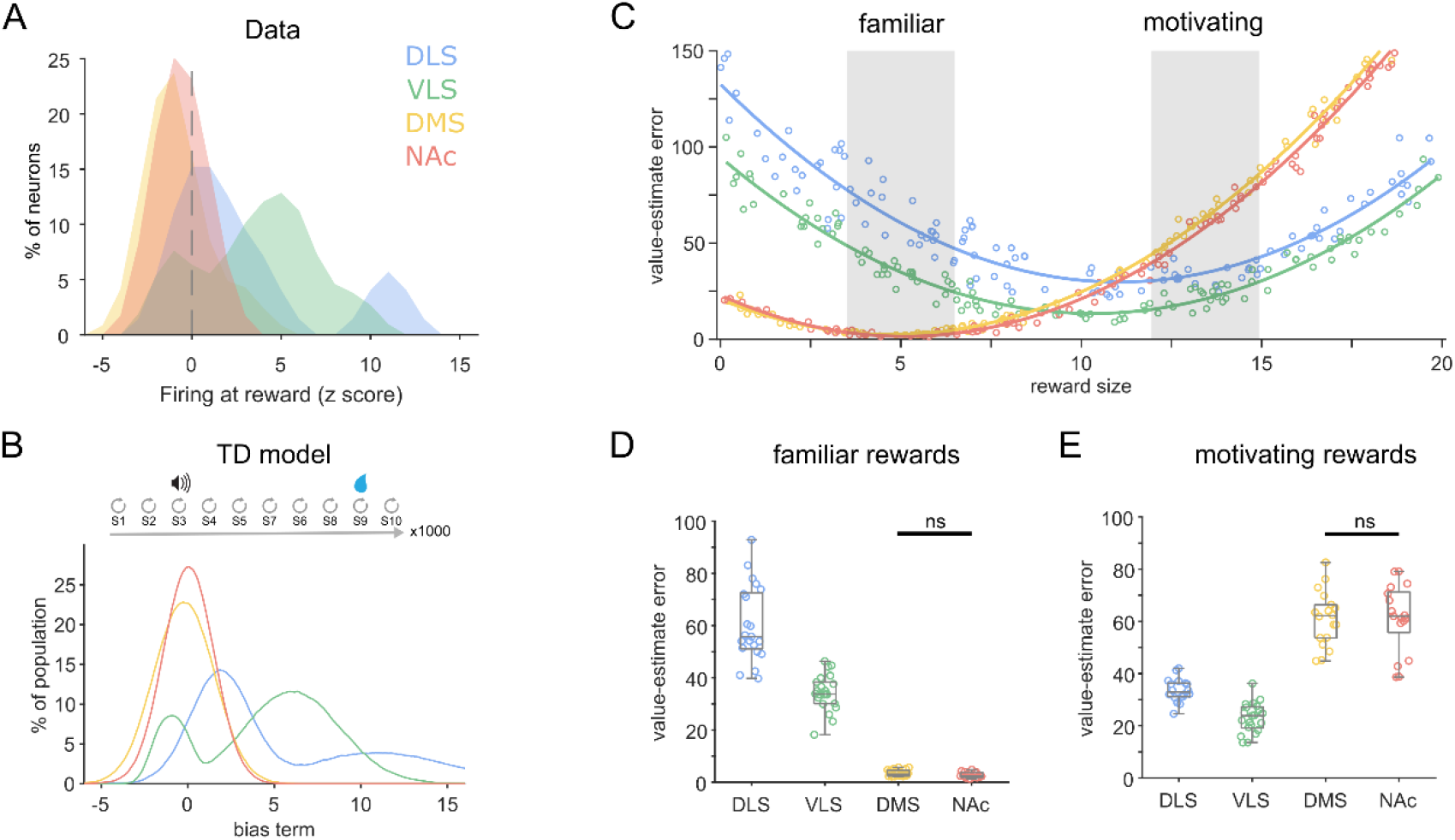
Different distributional coding of reward supports specialized roles in behavior. (A) Histogram of firing rate at reward for neurons in each putative projection-defined population. (B) Schematic (top) illustrating the temporal difference model where an agent transitions through states making state-value associations. Histogram (bottom) of the distribution of sampled bias terms (drawn from representations of the dopamine subpopulations in A) used to generate different projection-defined agents. (C) The difference between predicted and actual state values (mean squared error) for different numerical rewards. Grey shading indicates ranges used for familiar and motivating rewards. VLS-projecting agents exhibited significantly more accurate performance for all rewards > 50% of the reward range (or 5 standard deviations from the training mean). (D) Value-estimate errors of each agent to rewards in the familiar range (a range near the centre point of the training distribution). DLS vs VLS p < 0.001; DLS vs DMS or NAc p < 0.001; VLS vs DMS or NAc p < 0.001. Kruskal-Wallis one-way ANOVA on ranks with Dunnett post-hoc test. (E) Value-estimate errors of each agent to rewards in a motivating range (60-75% of the tested reward distribution). DLS vs VLS p < 0.001; DLS vs DMS or NAc p < 0.001; VLS vs DMS or NAc p < 0.001. Kruskal-Wallis one-way ANOVA on ranks with Dunnett post-hoc test. ns indicates no significant difference

## DISCUSSION

Here, we defined at millisecond resolution the behavior-related activity of individual, precisely localized, dopamine neurons and ascribed them to projection-defined cell populations. In doing so, we identified considerable heterogeneity in the encoding of reward-related signals by midbrain dopamine neurons. Heterogeneity was not well fit by anatomical subdivisions or spatial location, but could be better explained by grouping neurons according to the striatal regions they innervate. We show that individual dopamine neurons not only multiplex signals by encoding different combinations of egocentric and allocentric variables, but they also exhibit different magnitudes of encoding from the rest of the population. Our TD modelling predicts that such distributional coding not only maximizes the dynamic range of dopamine signals, but also tailors them to support the specialized roles of different striatal regions in learning and motivating behavior.

The role of dopamine neurons in predicting reward forms an important foundations for the understanding of how animals learn (Schultz, 2007). The dopamine signal has traditionally been considered to be uniform, broadcasting a common teaching signal across many brain circuits (Schultz, 2016a, 2016b). Instead, we find that such signals are far from uniform and our data suggest that different striatal regions receive specialized dopamine signals. Previous studies have identified that heterogeneity in dopamine neuron signaling can be parsed according to spatial localization in the midbrain (Dodson et al., 2016; Engelhard et al., 2019; Matsumoto and Hikosaka, 2009). Our initial analyses failed to observe homogenous responses segregated by location (Figure 3); however, when we considered neurons that putatively project to the same projection targets (Figure 5), we observed more homogeneous responses. The anatomical location of these projection-defined groups suggests that there are “hot spots” containing neurons projecting to the same region e.g. in parts of the medial substantia nigra pars compacta (SNCM) and the VTA (Figure 5D). Indeed these “hot spots” may explain why some studies observe more uniform responses when recording sites are more localized (Engelhard et al., 2019; Eshel et al., 2016).

Recent work has suggested that dopamine neurons not only encode RPE, but may also encode other task variables including movement onset and kinematics (Coddington and Dudman, 2018; Dodson et al., 2016; Engelhard et al., 2019; Howe and Dombeck, 2016; Hughes et al., 2020; da Silva et al., 2018). In addition to neurons encoding general body movements, we identified a number of neurons that encoded licking (Figures 1D, 1E and S1). It is not clear whether these signals represent a motor or perceptual response; in principle, firing at licking could signify the initiation of a tongue movement, the sensory properties of the reward, or something in between (Coddington and Dudman, 2018). This is further complicated by the possibility that there could simultaneously be a motor response in neurons projecting to DLS but an incentive response in those projecting to the NAc core; further work will be needed to disambiguate these possibilities. While many individual dopamine neurons encoded the cue, in contrast to previous studies (Flagel et al., 2011; Schultz et al., 1997), we did not observe a significant net response to the cue across the whole population (Figure 1C). One possible explanation for this is that we averaged neurons recorded throughout SNc and VTA; if some neuronal populations (e.g. the DMS-projecting group) decrease their firing at cue presentation, this could blunt the average response. However, even if these neurons were to be discounted, there is still not a large increase in firing (‘positive response’) to cue. It is unlikely to be due to inter-animal differences because for nearly every positive cue response we recoded, we also recorded other dopamine neurons in the same session with absent or negative cue responses. Another possibility is that the cue we used only had a modest volume (62 dB), which is considerably quieter than many commercial systems (which can be 75 – 86 dB). Dopamine neurons show larger responses to strong sensory stimuli, and diminished responses to cues predicting rewards with 100% reliability (Kutlu et al., 2021; de Lafuente and Romo, 2011). It is therefore possible that introducing louder cues or omitting rewards in some trials would unmask a larger increase in firing to the cue (Coddington and Dudman, 2018). However, our data suggest that a positive dopamine response at cue presentation is not necessary for task performance.

We identified combinations of marker expression and cell body location that could be used to delineate which striatal region a dopamine neuron is likely to target. This work complements studies investigating molecularly-identified dopamine neuron populations (Hook et al., 2018; La Manno et al., 2016; Pereira Luppi et al., 2021; Poulin et al., 2014, 2018, 2020; Saunders et al., 2018a; Tiklová et al., 2019; Wu et al., 2019) and provides additional confirmation of projection targets for some of these populations. While DLS- and NAc core-projecting populations exhibited “all or nothing” expression of three key markers, DMS- and VLS-projecting populations were less clear cut. This raises the possibility that there may be more than one population of dopamine neurons projecting to these regions. For example, in addition to the Sox6+ Aldh1a1-population projecting to VLS we identified, there could also be some Sox6-Aldh1a1+ dopamine neurons which are likely to be located more dorsally as Aldh1a1 expression decreases in ventral regions (Pereira Luppi et al., 2021; Poulin et al., 2018; Wu et al., 2019).

Our data suggest that dopamine neurons projecting to different regions have distinct firing patterns, an observation which is consistent with reports of differences in dopamine dynamics in different parts of striatum (Brown et al., 2011; Hamid et al., 2021; Heymann et al., 2019; de Jong et al., 2019; Lerner et al., 2015; Parker et al., 2016; Saddoris et al., 2015; Tsutsui-Kimura et al., 2020; Yuan et al., 2019). Perhaps the most striking difference between responses we observed was that the DMS-projecting group did not respond to reward; this finding is in agreement with some studies (Parker et al., 2016) but not others (Brown et al., 2011; Lerner et al., 2015). The fact that reward probability was deterministic in our experiment may help to reconcile these apparent discrepancies; a recent study compared dopaminergic axon terminal responses in DMS during fixed-vs variable-probability reward and observed a similar lack of response to fixed reward which was rescued as rewards became probabilistic (Tsutsui-Kimura et al., 2020). In contrast to DMS, we observed that the VLS-projecting population responded strongly to reward and that NAc core-projecting neurons responded during licking. Similar pronounced reward signals have previously been observed in dopamine neuron terminals within VLS, and delayed signals in medial regions which might be consistent with licking rather than reward (de Jong et al., 2019). Interestingly, aversive taste is reported to result in dopamine release preferentially in NAc core, suggesting a possible evaluative role for these licking-related signals (Yuan et al., 2019).

What are the implications of projection-selective-encoding? Because dopamine signals likely result in different outcomes depending on the target region (e.g. cue attraction vs movement invigoration) (Heymann et al., 2019; Saunders et al., 2018b), it follows that different striatal territories might receive distinct dopamine signals. Such specialized signals would permit flexibility and a wide dynamic range; for example, different regions might receive a common signal in one learning scenario for appetitive situations where approach is desired, but tailored signals in aversive scenarios where avoidance would be the appropriate behavior (de Jong et al., 2019; Lerner et al., 2015). We find that responses are tuned to different parts of the reward spectrum. For example, the positively skewed DLS- and VLS-projecting responses (Figure 6A) suggest populations of dopamine neurons which are tuned to better evaluate larger rewards. As such distributional coding within projection-defined populations may imbue further specialization (Dabney et al., 2020; Tsutsui-Kimura et al., 2020).

In conclusion, we find that even in simple learning paradigms, dopamine neurons represent multiple task variables in a heterogeneous manner. However, our data reveal an organizational logic where different striatal regions receive dopamine signals that are specialized to support different aspects of learning.

## AUTHOR CONTRIBUTIONS

Conceptualization, P.D.D; Methodology, P.D.D, A.K.K, and C.J.Y; Formal Analysis, R.A, A.K.K, C.J.Y, P.D.D, and G.E.P; Investigation, A.K.K, R.A, S.C, and G.E.P; Writing – Original Draft, P.D.D and R.A; Writing – Review & Editing, P.D.D, R.A, A.K.K, P.J.M, G.E.P, C.J.Y, R.P.C; Visualization, P.D.D; Supervision, P.D.D, P.J.M, and R.P.C; Funding Acquisition, P.D.D, P.J.M, and R.P.C.

## ACKNOWLEDGMENTS

We thank Mark Walton for helpful advice and Joe Pemberton and Dabal Pedamonti for feedback on an early version of this manuscript. We also thank Bristol and MRC BNDU animal services staff for expert technical assistance and the Wolfson Bioimaging Facility (University of Bristol) for microscopy support. This work was supported by BBSRC (BB/P006957/2 and BB/T013907/1 Awards to P.D.D), the MRC (Awards MC_UU_12024/2 and MC_UU_00003/5 to P.J.M), and a Monument Trust Discovery Award from Parkinson’s UK (J-1403 to P.D.D and P.J.M). A.K.K was supported by an MRC Doctoral Training Award (MC_ST_U13067) and C.J.Y and G.E.P were supported by the BBSRC South West Biosciences Doctoral Training Programme (BB/T008741/1 and BB/M009122/1).

## MATERIALS AND METHODS

### Experimental Animals

All experimental procedures on animals were conducted in accordance with the Animals (Scientific Procedures) Act, 1986 (United Kingdom) and approved by the animal welfare and ethical review boards at the University of Bristol and the University of Oxford. Unless otherwise stated, 3 – 4 month-old male C57Bl6/J mice (Charles River Laboratories) were used. Animals were maintained under a 12/12-h light/dark cycle, and experimental procedures were performed during the light phase of the cycle.

### Headpost implantation

For headpost implantation, mice were anesthetized using 1 – 2% (vol/vol) isoflurane. Mice were placed in a stereotaxic frame and on a homeothermic heating mat (Harvard Apparatus) to ensure stable body temperature. Corneal dehydration was prevented using carbomer liquid gel (Viscotears, Alcon) and mice were perioperatively injected with the analgesic buprenorphine (0.03 mg/kg s.c., Vetergesic, Bayern). A custom L-shaped headpost (0.7 – 0.8 g, stainless steel or aluminum) was attached to the skull using cyanoacrylate glue (Dodson et al., 2016). The 3 mm diameter window in the headpost-base was positioned above the substantia nigra of the right hemisphere (centered at AP −3 mm and ML +1.5 mm from bregma). A craniotomy for single-unit recordings was made within the window of the headpost either on the day of headpost implantation or 1 – 7 days prior to recording. Two stainless steel screws (0.8 mm diameter; Precision Technology Supplies) were implanted in the skull, one above the frontal cortex and a reference above the cerebellum of the left hemisphere. A coiled 0.23 mm diameter stainless-steel wire (AM Systems) was implanted between the layers of cervical muscle to record EMG activity (filtered at 0.3 – 0.5 kHz). Exposed skull, screws and EMG wire were covered with dental acrylic resin (Jet Denture Repair; Lang Dental). The craniotomy was sealed with fast set removable silicone rubber (Body Double; Smooth-On).

### Behavioral task

Animals were head-fixed using a custom headpost holder connected to a stereotaxic frame and positioned upon a custom-made treadmill where they could run, walk, or rest at will. Mice (N=37) were trained to associate an auditory cue with the delivery of a reward in a Pavlovian conditioning paradigm using a custom Arduino-based apparatus. Trials consisted of cue presentation (1 second, 4 kHz, 62 dB) delivered by a piezo speaker (535-8253, RS components), 1 second delay, followed by reward delivery (5 μl of 10% sucrose). Inter-trial interval (ITI) durations were randomly drawn from an exponential distribution with a flat hazard function to ensure equal distribution of expectation (4 – 10 s, median 5.4). Mice were either food (to > 85% of baseline weight) or water restricted (4 hours of *ad libitum* water per day after training/recording sessions). Animals were trained in daily sessions consisting of 100 rewards and all mice showed robust anticipatory licking to cue (median 5 days training prior to recording, IQR 3). Licking was monitored using a piezoelectric sensor (285-784, RS components). Movement and licking periods for single-unit recordings were determined off-line using cervical EMG and video recordings (30 frames/s). Lick-onset was defined as the first video frame with visually detectable lower jaw movement, lick-offset was defined as the first of a series of at least three subsequent video frames with no visually detectable jaw movement. Movement onset and offset were defined in the same way using body and limb movements.

### Electrophysiological recording and analysis

Extracellular single-unit recordings were made with borosilicate glass electrodes (tip diameter 1.0 – 1.5 μm, in situ resistance 10 – 25 MΩ; GC120F-10, Harvard Apparatus) filled with saline solution (0.5 M NaCl) containing Neurobiotin (1.5% w/v, Vector Laboratories). Sterile saline (0.9% w/v NaCl) was frequently applied around the craniotomy to prevent dehydration of the exposed cortex. Electrode signals were filtered at 0.3 – 5 kHz and amplified 1000 times (ELX-01MX and DPA-2FS, NPI Electronic Instruments). A Humbug (Quest Scientific) was used to eliminate mains noise at 50 Hz. For the recording, electrodes were lowered into the brain using a micromanipulator (IVM-1000; Scientifica). To avoid possible sampling bias, on-line criteria were applied to guide recordings of dopamine neurons (spike duration threshold-to-trough for bandpass-filtered spikes > 0.8 ms and firing rates < 20 spikes/s) (Janezic et al., 2013). Following recording, single neurons were juxtacellularly labeled with Neurobiotin (Dodson et al., 2016) to allow for their unambiguous identification and localization. At the end of the experiment, mice were given a lethal dose of pentobarbital and transcardially perfused with PBS followed by 4% w/v paraformaldehyde in 0.1 M phosphate buffer (PFA). Brains were placed in PFA overnight at 4°C and then stored in PBS containing 0.05% w/v sodium-azide.

In a subset of experiments (Figure 1, n=16 neurons), optotagging was performed instead of juxtacellular labelling. In these experiments, male DAT-IRES-Cre mice (n=5, B6.SJL-Slc6a3tm1.1(cre)Bkmn/J, Jackson Laboratories) were injected with AAV5-Ef1a-DIO-hChR2(E123T/T159C)-eYFP into the midbrain (~300 ul, AP −3.0, ML −1.0, DV −4.3), 4 – 6 weeks before headpost implantation to allow for robust viral expression. During recording, a 200 μm core (0.38 NA) fiber was placed inside the glass electrode. Laser pulses (5 ms, 1 Hz, 20 pulses, ~1mW) were used to stimulate neurons. A neuron was considered as opto-tagged (and therefore dopaminergic) if it fired spikes within 5 ms of >80% of the laser pulses.

All biopotentials were digitized online at 20 kHz using a Power 1401 mk3 analog-digital converter (Cambridge Electronic Design) and acquired using Spike2 software (version 7 or 10; Cambridge Electronic Design). Single-unit activity was isolated using template matching, principal component analysis and supervised clustering within Spike2 and data were exported to MATLAB (Mathworks). Firing activity of labeled neurons was normalized as z-scores and used to construct peri-stimulus time histograms (PSTH; bin size 40 ms, smoothed with a 5-point Gaussian filter half-width 70 ms) using a baseline of 1 second preceding cue onset. We considered firing at reward to be the average z score of the four bins starting 40 ms after reward delivery. To analyze which factors accounted best for changes in firing of individual midbrain dopamine neurons recorded in the conditioning experiment, a generalized linear regression model (glmfit function, MATLAB) was used to obtain a least-squares fit of the selected predictors to the recording data. Predictors were defined as cue, reward, licking, and movement. Regression coefficients were estimated for each labeled neuron in 200 ms windows aligned to cue onset. An individual cell was considered responsive to one of the four variables if the corresponding p-value was < 0.01. Because periods of licking often occurred soon after reward delivery, we confirmed that the firing of neurons classified as ‘licking’ was time locked to lick episodes but not reward delivery (Figure S1). To analyze dopamine neuron firing properties, we used a coefficient of variability of interspike intervals (CV2) to examine firing regularity (Holt et al., 1996; Janezic et al., 2013) and robust Gaussian surprise (Ko et al., 2012; Sloan et al., 2016) to detect bursts of at least three spikes with significantly shorter ISI’s than the population of spike trains.

The Shapiro-Wilk test and the Levene test were used to judge whether data sets were normally distributed with homogeneous variances (p < 0.05 to reject). For normally distributed data, a one-way ANOVA was used. If data failed normality tests, Mann-Whitney rank sum or Kruskal-Wallis one-way ANOVA on ranks with Dunnett’s post-hoc method for multiple comparisons were used (MATLAB, Mathworks). Significance for statistical tests was set at p < 0.05. Boxplots display first quartile, median and third quartile.

### Retrograde tracer injections

Mice were prepared for surgery as previously described; a craniotomy was performed above the target region and a calibrated glass micropipette (708707; Blaubrand IntraMark) with a tip diameter of ~25μm was lowered to the appropriate target; NAc core (n = 7 mice; AP +1.0, ML +1.0, DV −4.3), DLS (n = 6 mice; AP +1.1, ML +1.8, DV −3), DMS (n = 5 mice; AP +1.0, ML +1.2, DV −2.8), VLS (n = 5 mice; AP +1.0, ML +1.8, DV −4.2). 30 – 150 nl cholera toxin subunit b (CTB; 0.5% w/v; C9903; Sigma-Aldrich) was manually injected at a rate of ~50 nl/min and pipettes were left in place for 5 – 10 minutes after injection. 9 – 13 days after tracer injection, mice were given a lethal dose of anesthetic and transcardially perfused as previously described. In a minority of experiments (n = 8), we injected CTB into dorsomedial striatum prior to recording, to verify that recorded neurons projected to the putatively assigned target; in these experiments we recorded and juxtacellularly labeled two SNCM neurons, both of which were CTB positive.

### Immunohistochemistry

50 μm coronal sections were cut from the midbrain on a vibrating-blade microtome (VT1000S; Leica Microsystems or DTK-1000, DSK). To confirm the location and neurochemical identity of recorded and juxtacellularly-labeled neurons, sections were incubated for 4 h at room temperature in PBS with 0.3% (vol/vol) Triton X-100 (Sigma) containing Cy3-conjugated streptavidin (1:1000) (GE Healthcare). To probe expression of different molecular markers in labeled neurons, a two-step procedure was applied, sections were incubated overnight in PBS-Triton with mouse anti-Tyrosine Hydroxylase (TH, 1:1000, T2928, Sigma-Aldrich) or chicken anti-TH (1:500, ab76442, Abcam); guinea pig anti-Sox6 (1:1000, gift from M. Wegner, Friedrich-Alexander University Erlangen-Nuremberg; (Stolt et al., 2006)) or rabbit anti-Sox6 (1:500, ab30455, Abcam). Sections were washed in PBS, and then incubated in PBS-Triton for > 4 hours with AMCA-conjugated secondary antibodies (donkey anti-mouse IgG, 1:500; 715-155-150 or donkey anti-chicken IgG, 1:500, 703-155-155; Jackson ImmunoResearch) or Brilliant Violet 421-conjugated secondaries (donkey anti-chicken IgG 1:500, 703-675-155, Jackson ImmunoResearch) to visualize immunoreactivity for TH, and AlexaFluor 647- or Cy5-conjugated secondary antibody to visualize immunoreactivity for Sox6 (A647: donkey anti-guinea pig IgG, 1:500, 706-605-148, Jackson ImmunoResearch; Cy5: donkey anti-rabbit IgG, 1:500, 711-175-152, Jackson ImmunoResearch). After imaging, the second step consisted of incubating overnight in PBS-Triton with rabbit anti-Aldh1a1 (1:500, HPA002123, Sigma Merck) and goat anti-calbindin (1:500, sc7691; Santa Cruz) or mouse anti-calbindin(1:500, CB300, Swant), washing in PBS and then incubating overnight at room temperature in PBS-Triton with AlexaFluor 647- or Cy5-conjugated secondary antibodies (the fluorophore used in the previous step to visualize Sox6) to visualize immunoreactivity for Aldh1a1 (AF647: donkey anti-rabbit IgG, 1:500, 711-605-152, Jackson ImmunoResearch; Cy5: donkey anti-rabbit IgG, 1:1000, 711-175-152, Jackson ImmunoResearch) and AlexaFluor 488-conjugated secondary antibodies for Calbindin (donkey anti-goat IgG, 1:500, A11055, Life Technologies; donkey anti-mouse IgG, 1:500, 715-545-150, Jackson ImmunoResearch). This way, we were able to clearly visualize immunoreactivity for Sox6 (nuclear) and Aldh1a1 (cytoplasmic) using the same fluorescence channel. Borders of VTA and SNc were delineated using Aldh1a1 and calbindin immunofluorescence (Fu et al., 2012).

For retrograde tracing, a combinatorial approach with partial overlap was used for this immunostaining protocol, so that TH and CTB immunoreactivity was tested in all samples, but different series from the same animal were tested for Sox6, Aldh1a1, and calbindin. Sections were incubated overnight at room temperature in PBS-Triton with chicken anti-TH (1:250/500, ab76442, Abcam), mouse anti-CTB (1:500, ab35988, Abcam) or goat anti-CTB (1:5000), #703, List Biological Labs), rabbit anti-calbindin (1:1000, CB38, Swant) or goat anti-calbindin (1:500, sc7691, Santa Cruz) or mouse anti-calbindin (1:500, CB300, Swant), rabbit anti-Sox6 (1:4000, ab30455, Abcam), rabbit anti-Aldh1a1 (1:500, HPA002123, Sigma Merck). Sections were washed in PBS and then incubated for > 4 h at room temperature in PBS-Triton and secondary antibodies. To visualize immunoreactivity for TH, AMCA- or Brilliant Violet 421-conjugated secondary antbodies were used (AMCA: donkey anti-chicken IgG, 1:500, 703-155-155, Jackson ImmunoResearch; BV421: donkey anti-chicken IgG 1:500, 703-675-155, Jackson ImmunoResearch). CTB was visualized using Cy3- or AlexaFluor 488-conjugated secondaries (Cy3: donkey anti-mouse IgG, 1:500, 715-165-151, Jackson ImmunoResearch; AF488: donkey anti-goat IgG, 1:500, 705-545-147, Jackson ImmunoResearch). Aldh1a1 or Sox6 immunoreactivity was visualized using Cy5- or AlexaFluor 647-conjugated secondaries (Cy5: donkey anti-rabbit IgG, 1:1000, 711-175-152, Jackson ImmunoResearch; AF647: donkey anti-rabbit IgG, 1:500, 711-605-152, Jackson ImmunoResearch) and Calbindin with Cy3- or AlexaFluor 488-conjugated secondaries (Cy3: donkey anti-mouse IgG, 1:500, 715-165-150, Jackson ImmunoResearch; AF488: donkey anti-mouse IgG, 1:500, A-21202, Life Technologies).

### Microscopy and cell counting

Example images were acquired using a confocal laser-scanning microscope (20x objective, LSM710; Carl Zeiss, or SP8; Leica). Images for cell counting were acquired on an epifluorescence microscope (DMI6000; Leica, or AxioImage.M2; Carl Zeiss) equipped with a 20x objective. Images of the dopaminergic midbrain were acquired as a series of 21 tiles (7x,3y). Sections containing CTB positive SNc and VTA neurons with a clearly defined nucleus were counted using the ‘cell counter’ plugin on ImageJ, Fiji version 1.53q (Schindelin et al., 2012) or Stereo investigator software 9.0 (MBF Bioscience). During counting, the experimenter was blind to the region targeted with CTB. To obtain percentages of midbrain dopamine neurons expressing a particular marker, counts were collapsed across sections, then divided by the number of neurons positive for both CTB and TH in each sample. Every marker-combination was counted in a minimum of three animals per striatal region. CTB injection sites in striatum were represented as honeycomb plots; a tessellated hexagonal structure was superimposed onto each image, then hexagons that were >80% by CTB immunoreactivity were coloured red at 100% opacity, opacity of hexagons that included 50 – 80% CTB immunopositivity was set at 50%. Images from each animal were superimposed and opacity was normalised.

### Computational modelling

For each observed state, our Temporal Difference (TD) algorithm (Schultz et al., 1997; Sutton, 1988) produced a series of value predictions (*V_t,i_*). Each of these Value predictions represents a single neuron (i, at time t). TD error (*δ_t,i_*) was calculated by comparing a neuron’s existing state-value prediction, with a bootstrapped (predicted) estimate of the state’s value (*V*_*t*+1_,_*i*_):

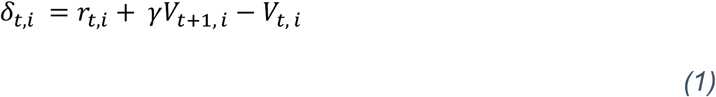

where (*r_t,i_*) is the reward and *γ* the discount factor. Value predictions were updated according to the following update rule:

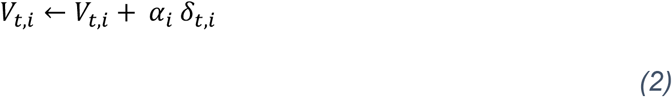

Where *α_i_* is the learning rate applied to each neuron (*α_i_* ~ U(0,1)). To model distributional encoding as a product of varied firing at reward, a bias term (*φ_i_*) is added to *δ_t_* which is drawn from the activity distributions of each dopamine neuron subpopulation. These subpopulations were approximated using kernel density estimation, thereby we propose that different populations encode RPE differently.

To test these algorithms, we created a ‘Pavlovian conditioning’ environment in which agents deterministically transition between states. At one such state, the agent receives a conditioned stimulus; after a delay (representing the agent transitioning between five states), the agent received a numerical reward randomly selected from a gaussian distribution (*r* ~ *N*(5,1) Trained agents have a ‘value distribution’ associated with each state in its environment; produced from the distribution of state-error associations across simulated neurons.

After training, agents were tested with different rewards. Agents were tested on a wider range (r ~ U (0,20)) of familiar (i.e. ~5) and larger rewards. To determine how accurate each agent was at estimating value, we calculated the mean squared error (MSE) between actual (*Y*) and predicted (*Ŷ*) value, for each cell at the rewarded state:

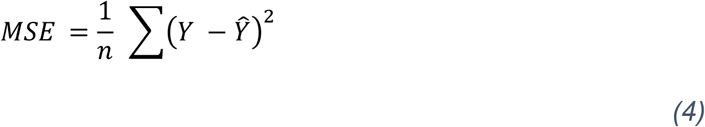

Note that predicted value (*Ŷ*) should approximate the reward received during testing.

## SUPPLEMENTAL INFORMATION

**Supplementary Figure 1.**
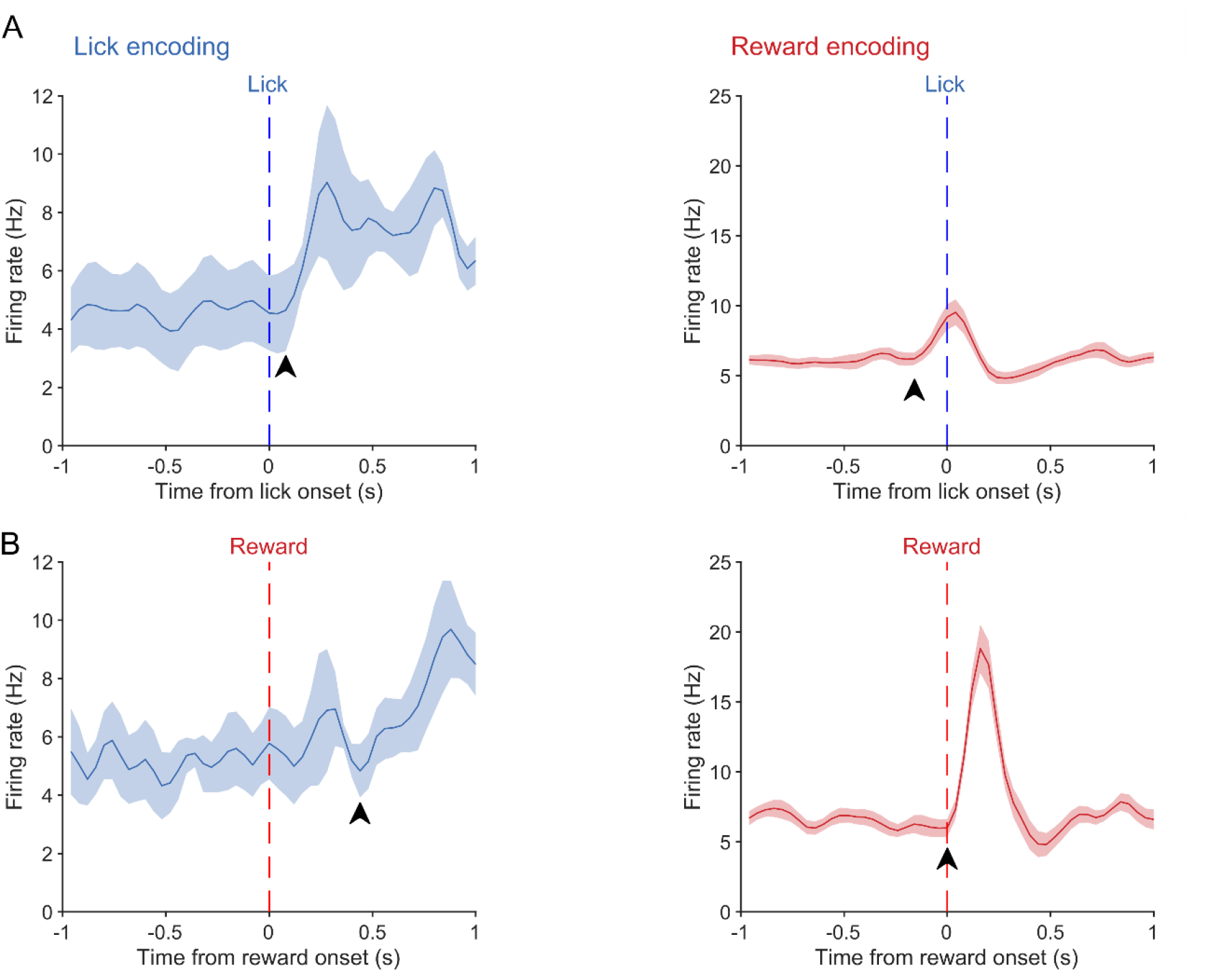
Increases in firing of neurons encoding licking are better matched to lick onset than reward. (A) The firing rate of neurons that significantly (and exclusively) encoded licking (n = 5) increases (arrowhead) just after lick onset (blue dashed line). Increases in the firing of neurons exclusively encoding reward (n = 20) precede lick onset. (B) Increases in the firing rate of neurons encoding reward (right) are well timed with reward delivery (red dashed line) unlike increases in firing rate in neurons that encoded licking (left)

**Supplementary Figure 2.**
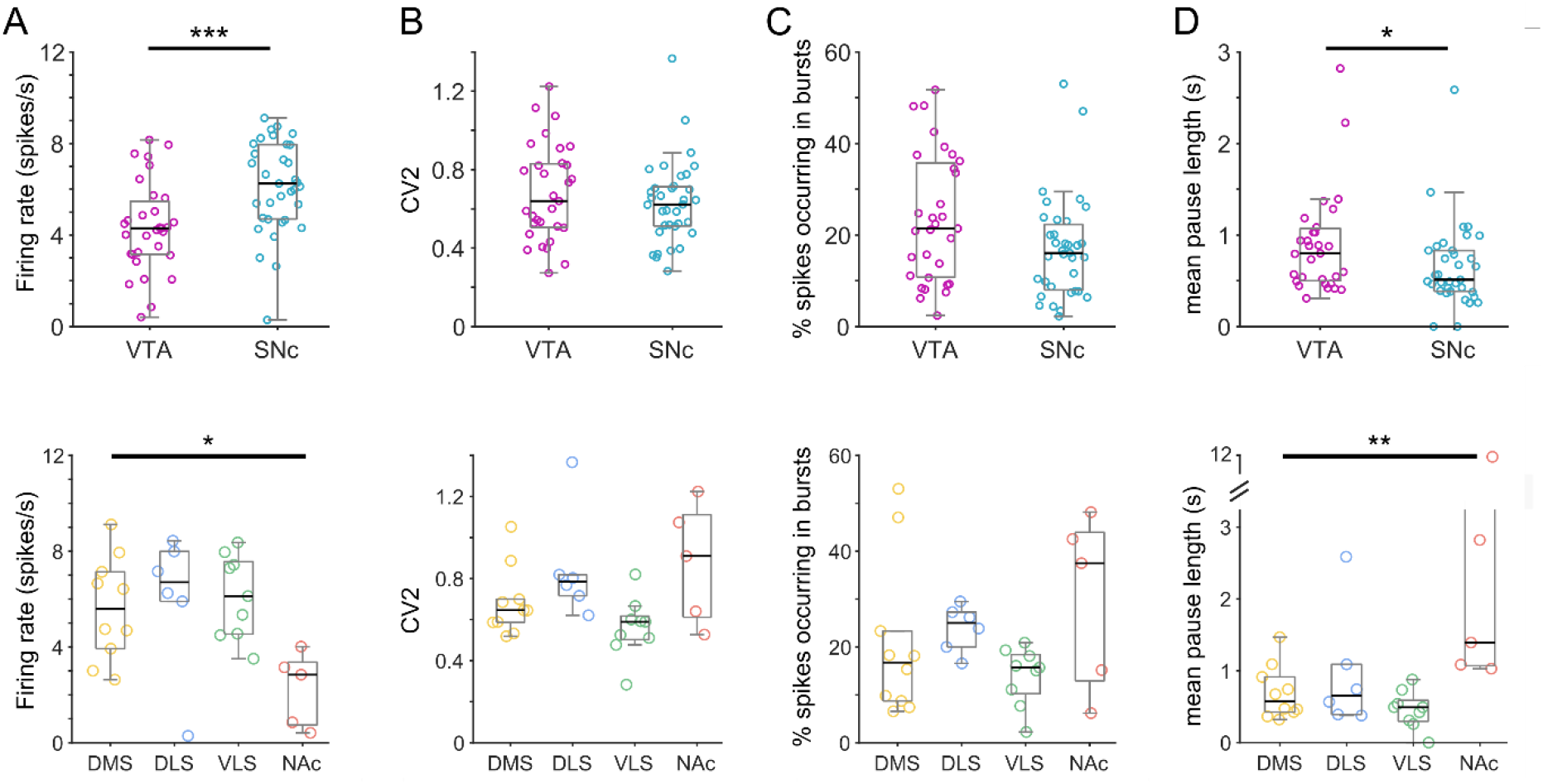
Distinct firing properties of dopamine neuron populations. (A) Firing rate was significantly lower in VTA neurons than SNc neurons (Mann-Whitney rank sum) and putative NAc core projecting neurons fired significantly slower than all other populations (Kruskal-Wallis one way ANOVA on ranks). There were no significant differences between DMS, DLS and VLS. (B) Firing regularity (CV2) was not significantly different between VTA and SNc neurons (Mann-Whitney rank sum) nor any of the projection-defined groups (Kruskal-Wallis one way ANOVA on ranks). (C) The proportion of spikes fired as bursts were not significantly different between VTA and SNc neurons (Mann-Whitney rank sum) nor any of the projection-defined groups (Kruskal-Wallis one way ANOVA on ranks). (D) The mean duration of pauses was significantly longer for VTA than SNc neurons (Mann-Whitney rank sum). Similarly, putative NAc core projecting neurons exhibited longer pauses in firing than other populations (Kruskal-Wallis one way ANOVA on ranks). * p < 0.05, ** p < 0.01, *** p < 0.001

